# The human UDP-galactose 4’-epimerase (GALE) is required for cell surface glycome structure and function

**DOI:** 10.1101/646794

**Authors:** Alex Broussard, Alyssa Florwick, Chelsea Desbiens, Nicole Nischan, Corrina Robertson, Ziqiang Guan, Jennifer J. Kohler, Lance Wells, Michael Boyce

## Abstract

Glycan biosynthesis relies on nucleotide-sugars (NS), abundant metabolites that serve as monosaccharide donors for glycosyltransferases. *In vivo*, signal-dependent fluctuations in NS levels are required to maintain normal cell physiology and are dysregulated in disease, but how mammalian cells regulate NS levels and pathway flux remains largely uncharacterized. To address this knowledge gap, we examined uridine diphosphate (UDP)-galactose 4’-epimerase (GALE), which interconverts two pairs of essential NSs. GALE deletion in human cells triggered major imbalances in its substrate NSs and consequent dramatic changes in glycolipids and glycoproteins, including a subset of integrins and the Fas death receptor. NS dysregulation also directly impacted cell signaling, as GALE^−/−^ cells exhibit Fas hypoglycosylation and hypersensitivity to Fas ligand-induced apoptosis. Our results reveal a new role for GALE-mediated NS regulation in supporting death receptor signaling and may have implications for the molecular etiology of illnesses characterized by NS imbalances, including galactosemia and metabolic syndrome.

---

Glycosylation – the enzymatic attachment of carbohydrates to proteins, lipids and other biomolecules – is an abundant and conserved modification across all clades of life (1). In mammals, glycosylation influences nearly every cell biological process, including protein quality control and secretion, adhesion and migration and host-pathogen interactions (2–4). Consistent with this central role in mammalian physiology, aberrant glycosylation contributes to the pathology of myriad human diseases, such as developmental defects, diabetes, obesity and metabolic syndrome, cancer, neurodegeneration and atherosclerosis (5–13).

Virtually all glycoconjugates are assembled from nucleotide-sugars (NSs), metabolites that donate “activated” monosaccharides to glycosyltransferases (2). In recent years, several groups observed that specific stimuli or signaling events, such as feeding or ischemic stress, trigger increased NS biosynthesis in mammalian cells, likely to facilitate protein secretion and support the remodeling of cell-surface glycans (14, 15). Many glycosyltransferases are sensitive to NS concentrations, such that changes in NS levels affect not only bulk levels of glycosylation but also specific glycosyltransferase substrate choice (16–21). These observations highlight the critical role of NS regulation in shaping downstream glycoconjugate biosynthesis and function. However, while the biochemistry of NS biosynthetic enzymes is well understood, little is known about how cells regulate flux through NS metabolic pathways in response to signals or disease states. Furthermore, the impact of NS fluctuations on key glycosylation pathways and downstream cellular phenotypes is poorly understood, representing a major knowledge gap in cell biology.

As a first step toward understanding the mechanisms and functions of human NS regulation, we focused on UDP-galactose 4’-epimerase (GALE) as a model enzyme. Mammalian GALE interconverts two pairs of substrates: the hexose NSs UDP-glucose (UDP-Glc)/UDP-galactose (UDP-Gal), and the corresponding *N*-acetylhexosamine NSs UDP-*N*-acetylglucosamine (UDP-GlcNAc) and UDP-*N*-acetylgalactosamine (UDP-GalNAc) (Fig. 1A) (22–24). Through these reversible epimerizations, human GALE balances the pools of four NSs essential for the biosynthesis of thousands of glycoproteins and glycolipids (2, 23, 24). Interestingly, however, GALE is not absolutely required for the biosynthesis of any of its four substrates, each of which could hypothetically be derived from independent salvage or *de novo* metabolic routes in nutrient-replete cells (2, 22). Because it acts only by interconverting existing NSs, GALE is an excellent model enzyme to study the role of dynamically balancing NS pools in cell physiology. Furthermore, GALE is significant to human health, both because GALE operates in the liver and hypothalamic neurons of healthy mammals to regulate Glc metabolism and satiety after feeding (14, 25, 26), and because partial loss-of-function GALE mutations cause a subtype of the congenital disease galactosemia and a rare form of thrombocytopenia (27–29).

**Figure 1.**
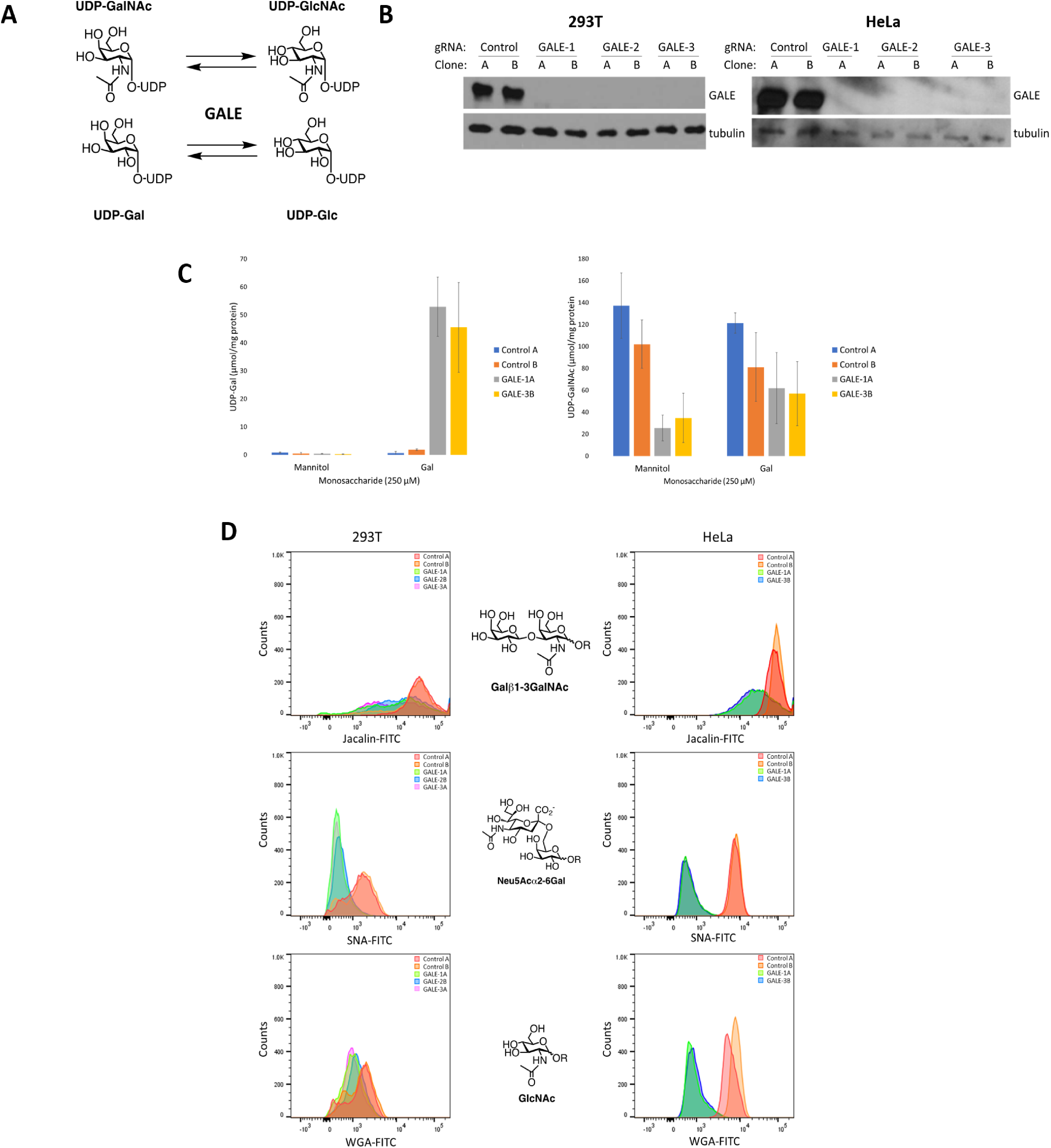
Human GALE is required for NS metabolism and glycoconjugate biosynthesis. (A) Human GALE epimerizes two pairs of NSs, UDP-Gal/UDP-Glc and UDP-GalNAc/UDP-GlcNAc. (B) GALE was deleted using CRISPR/Cas9 methods and one of three GALE-targeting sgRNAs (1-3) in 293T (left) and HeLa (right) cells. Single-cell derived clones (denoted A or B) were lysed and analyzed by Western blot. (C) Control and GALE^−/−^ HeLa cells were treated with 250 μM Gal or mannitol (osmolarity control) for 72 hours and UDP-Gal and UDP-GalNAc were quantified by high performance anion exchange chromatography (HPAEC). N=3 biological replicates. Error bars represent standard error of the mean (SEM). See also Figure S1. (D) Control and GALE^−/−^ 293T (left) and HeLa (right) cells were stained with fluorescently (FITC) tagged jacalin, SNA or WGA lectins and 10,000 cells from each sample were analyzed by flow cytometry. The major glycan ligand of each lectin is indicated between the corresponding panels.

To characterize the role of GALE in cell physiology, we used CRISPR/Cas9 methods to create GALE^−/−^ human cell systems. Our results reveal that GALE is required to maintain NS levels and to biosynthesize a wide range of glycoproteins and glycolipids, even under nutrient-replete conditions. In particular, we show that GALE is essential for the N-glycosylation of several cell-surface proteins, including a subset of integrin cell adhesion molecules and the apoptotic death receptor Fas. Moreover, GALE deletion results in Fas hypoglycosylation and hypersensitivity to Fas ligand (FasL)-induced cell death, highlighting a previously unknown function of NS metabolism in apoptotic pathways. Our results reveal a requirement for human GALE in supporting glycoconjugate biosynthesis and cell surface signaling, and establish loss-of-function culture systems as a powerful tool for dissecting the role of NS regulation in human cell biology.

## Results

### Human GALE is required to balance NS levels

GALE is the final enzyme in the Leloir pathway, a highly conserved metabolic route for the assimilation of Gal (Fig. 1A) (30). To determine the role of GALE in NS metabolism and glycoconjugate biosynthesis, we used CRISPR/Cas9 methods to construct multiple single cell-derived GALE^−/−^ clones from human cell lines and confirmed successful ablation of GALE protein (Fig. 1B). Based on prior studies of the Leloir pathway in human patients and experimental model systems, we hypothesized that GALE^−/−^ cells might display NS imbalances under standard culture conditions and/or in the presence of supplementary Gal (31, 32). In particular, Gal consumption is closely tied to adverse symptoms in galactosemic patients and laboratory models (33, 34). Therefore, we sought to determine the impact of Gal on viability and NS metabolism in our cell models. Gal supplementation did not impair cell viability in control or GALE^−/−^ cells (Fig. S1). However, GALE^−/−^ cells accumulated high levels of UDP-Gal in the presence of supplementary Gal, whereas control cells showed only a modest increase (Fig. 1C). Interestingly, GALE^−/−^ cells also exhibited reduced basal levels of UDP-GalNAc, which was partially rescued by Gal addition (Fig. 1C). We concluded that GALE is required to maintain normal NS levels in nutrient-replete human cells in the absence or presence of supplementary Gal.

### GALE is required for glycoprotein and glycolipid biosynthesis

Given the major NS imbalances observed in GALE^−/−^ cells (Fig. 1C), we hypothesized that they might display defects in the biosynthesis of glycoproteins and glycolipids containing Gal or GalNAc, such as mucin-type O-glycoproteins and gangliosides (35–37). Furthermore, absence or reduction of Gal/GalNAc moieties is predicted to disrupt terminal glycan structures, such as sialic acids, which are typically added to Gal or GalNAc residues of mature glycans (38). Flow cytometry assays with jacalin, a lectin that binds the Galβ1-3GalNAc Thomsen-Friedenreich antigen core of mucin-type glycoproteins (39), wheat germ agglutinin, which binds terminal GlcNAc (40), and *Sambucus nigra* lectin (SNA), which binds sialic acids (41), indicated that GALE^−/−^ cells indeed have reduced levels of each species, as compared to control cells (Fig. 1D). Consistent with these observations, monosaccharide composition analysis revealed a substantial decrease in total sialic acid, Gal and GalNAc levels in glycans isolated from GALE^−/−^ cells, compared to controls (Fig. 2A).

**Figure 2.**
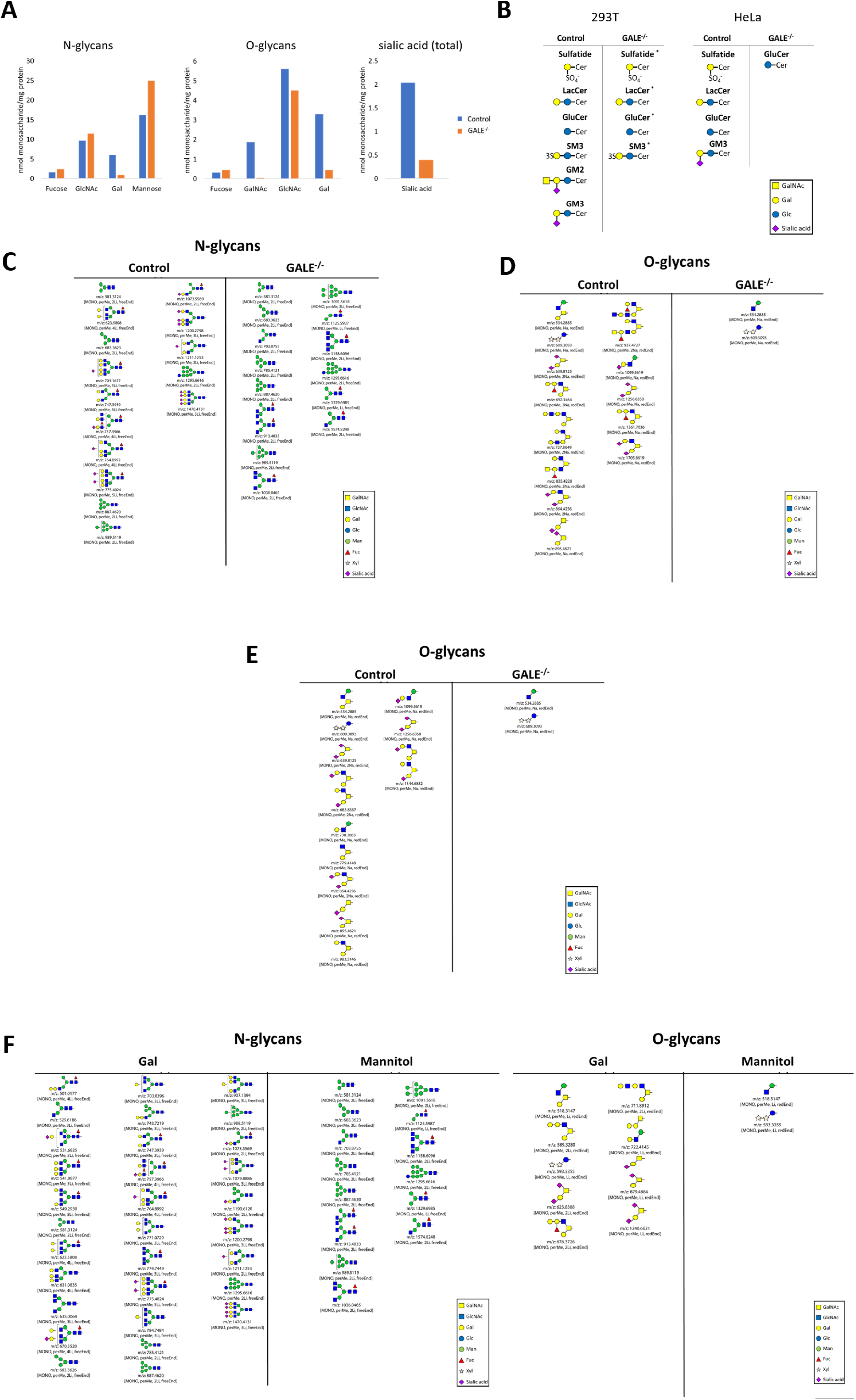
GALE deletion triggers global changes in cell surface glycoproteins and glycolipids. (A) Monosaccharide composition analysis of N- and O-linked glycans from control and GALE^−/−^ HeLa cells. (B) Summary of glycolipid species identified in control and GALE^−/−^ 293T (left) and HeLa (right) cells via LC-MS. Asterisks indicate glycolipid species dramatically reduced in GALE^−/−^ cells. See also Figure S2. (C) Summary of O-glycan species identified in control and GALE^−/−^ 293T cells via LC-MS. m/z values are given for each species. See also Figure S3. (D) Summary of O-glycan species identified in control and GALE^−/−^ HeLa cells via LC-MS. m/z values are given for each species. Xyl, xylose. See also Figure S3. (E) Summary of N-glycan species identified in control and GALE^−/−^ HeLa cells via LC-MS. m/z values are given for each species. Man, mannose; Fuc, fucose; MONO, monoisotopic mass; perme, permethylated; xLi or xNa, indicates lithium or sodium adduct with x ions; redend, reducing end; freeend, nonreducing end. See also Figure S4. (F) Summary of N-(left) and O-glycan (right) species identified by LC-MS in GALE^−/−^ HeLa cells treated with 250 µM Gal or mannitol (osmolarity control) for 72 hours. m/z values are given for each species. See also Figure S4.

We next profiled global glycolipids and protein N- and O-linked glycans via liquid chromatography/mass spectrometry (LC-MS). GALE^−/−^ and control cells bore similar levels of many lipids and glycolipid precursors, such as phosphatidylinositol, ceramide, and cardiolipin (Fig. S2). However, levels of Gal/GalNAc-containing glycolipids, such as sulfatides and several gangliosides, were greatly reduced or undetectable in GALE^−/−^ cells, compared to controls (Figs. 2B and S2). These results demonstrate the importance of GALE in maintaining the glycolipidome. In parallel, we analyzed N- and O-linked glycans from cell surface glycoproteins. Consistent with our lectin staining results, we observed striking deficiencies in both mucin-type O-glycoproteins and Gal/GalNAc-containing N-linked glycan structures in GALE^−/−^ cells (Figs. 2C-E, S3 and S4). Gal-responsive defects in glycoprotein synthesis have been observed in galactosemia subtypes caused by mutations in other Leloir pathway enzymes (42), but relatively little is known about how loss of GALE impacts glycan synthesis in the presence of Gal. To test the importance of GALE in this context, we analyzed N- and O-linked glycoproteins and showed that the glycan profiles of GALE^−/−^ cells treated with Gal (but not those treated with mannitol, a non-metabolizable osmolyte control) closely resembled the glycans of controls cells (Figs. 2C-F, S3 and S4). We concluded that human GALE is required to support the biosynthesis of a broad range of glycoproteins and glycolipids.

### GALE is required for cell-surface receptor glycosylation and function

We reasoned that altered glycosylation in GALE^−/−^ cells might impact signaling through mis-glycosylated receptors. Consistent with this hypothesis, we observed substantial molecular weight shifts in several specific glycoproteins in GALE^−/−^ cells, compared to controls, including death receptors and integrins, major mediators of cell-matrix adhesion (Figs. 3A, 3B and S5). Gal supplementation suppressed these molecular weight changes in GALE^−/−^ cells, indicating that they are caused by hypoglycosylation in the absence of GALE, rather than an off-target or indirect effect of CRISPR manipulation (Figs. 3A and S5). Interestingly, although most integrins are glycoproteins (43, 44), only a subset was impacted by GALE deletion, suggesting specific roles for GALE activity in the biosynthesis of particular glycoconjugates (Fig. S5). Our LC-MS studies indicated a relatively modest role for GALE in global N-glycan biosynthesis, as compared to O-glycans (Figs. 2C-E). However, several key surface receptors, including Fas and integrin β_1_, exhibited aberrant N-glycoforms, as treatment with the glycosidase PNGase-F, which cleaves N-glycans (45) restored these proteins to their predicted molecular weights in both control and GALE^−/−^ cells (Fig. 3B).

**Figure 3.**
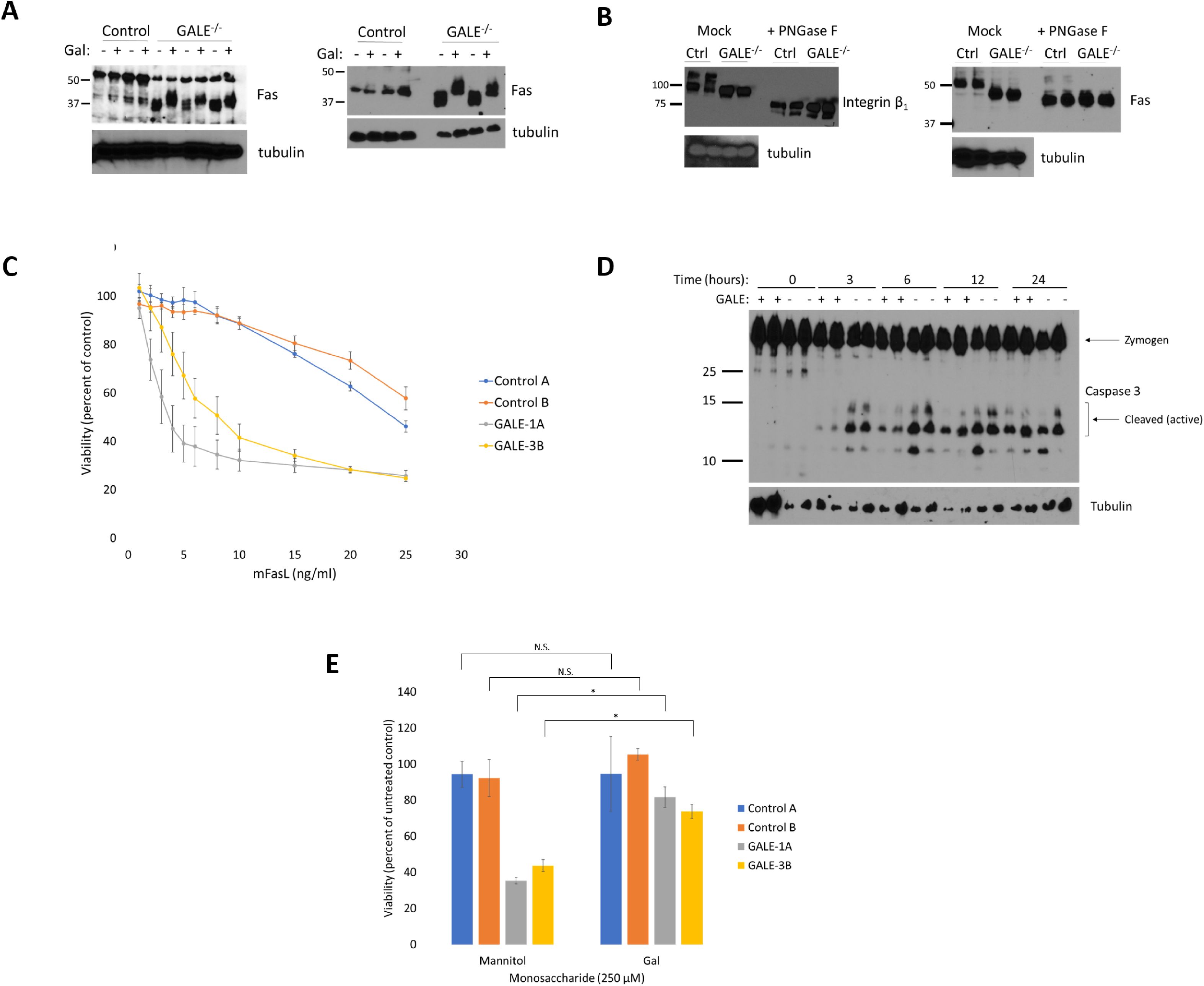
GALE is required for Fas glycosylation and function. (A) Control and GALE^−/−^ 293T (left) and HeLa (right) cells were treated with 250 µM Gal or mannitol for 72 hours and lysates were analyzed by Western blot. Each pair of lanes (+/-Gal) represents a distinct clone. (B) Lysates from control and GALE^−/−^ HeLa cells were treated with PNGase-F and analyzed by Western blot, highlighting GALE-dependent N-glycosylation in integrin β_1_ (left) and Fas (right). Within treatment groups (+/-PNGase-F), each lane represents a distinct clone. See also Figure S5. (C) Control and GALE^−/−^ HeLa clones were treated with the indicated concentrations of FasL for 24 hours and cell viability was measured by MTS assay. N=3 biological replicates. Error bars represent SEM. *p* < 0.05 for all FasL concentrations ≥ 6 ng/ml, comparing control to GALE^−/−^ (two-way ANOVA, post hoc one-way ANOVA). See also Figures S6 and S7. (D) Control and GALE^−/−^ HeLa clones were treated with 2.5 ng/ml FasL for the indicated times and lysates were analyzed by Western blot. See also Figure S6.

To test the function of GALE-dependent glycosylation, we focused on the death receptor Fas as a model glycoprotein. Fas trimerization and activation by FasL trigger the formation of the Death-Inducing Signaling Complex (DISC), which recruits and activates upstream caspases, ultimately leading to downstream caspase-3 activation and apoptotic death (46, 47). No role for GALE has been reported previously for death receptor signaling, but prior work demonstrated that changes in Fas sialylation affect DISC formation and the apoptotic cascade (48). Therefore, we hypothesized that Fas hypoglycosylation might affect cell death signaling in GALE^−/−^ cells.

Neither GALE deletion nor Gal treatment altered the cell-surface expression of Fas (Fig. S6). Interestingly, however, GALE^−/−^ cells were significantly sensitized to FasL-induced killing (Fig. 3C). This result reflects a specific effect on FasL-induced death, rather than a general susceptibility to apoptosis, because no analogous effect was observed when cells were treated with the broad-spectrum serine/threonine kinase inhibitor staurosporine (Fig. S7). Next, to confirm the mechanism of sensitization to FasL in GALE^−/−^ cells, we examined the apoptotic executioner caspase-3 (46). Compared to controls, GALE^−/−^ cells exhibited increased caspase-3 cleavage (indicative of its activation) at early time-points after FasL exposure (Fig. 3D). Finally, we examined the specific GALE-dependent glycan determinants that influence Fas signaling. Prior work by Bellis and colleagues demonstrated that upregulation of the ST6Gal-I sialyltransferase in tumor cells potentiates Fas sialylation, reducing its responsiveness to ligand activation (48). Based on these reports and our monosaccharide composition and glycoprotein profiling data (Figs. 2A, 2E, 2F), we hypothesized that the hypersensitivity of GALE^−/−^ cells to FasL was caused by Fas hyposialylation. Consistent with this idea, Gal supplementation reversed the sensitivity to FasL in GALE^−/−^ cells, indicating a GALE-dependent causal link between receptor glycosylation (Figs. 3A and 3B) and activity (Figs. 3E and S8). Taken together, these results demonstrate that GALE is required to support the biosynthesis of sialylated N-glycans on Fas and other surface proteins, and loss of GALE function dysregulates glycoprotein receptor signaling.

## Discussion

The regulation of human NS levels has profound implications for both normal physiology and disease, yet it remains poorly understood. Our results shed new light on the role of GALE, a model NS metabolic enzyme, in the biosynthesis and function of a wide range of glycoconjugates.

### GALE governs human NS levels

GALE^−/−^ cells have greatly reduced levels of UDP-GalNAc and UDP-Gal, compared to controls (Fig. 1C). Based on these observations, we propose that GALE is required for the biosynthesis and balancing of these key NS, despite the presence of free trace Gal and Gal/GalNAc-containing glycoproteins in serum, which could hypothetically be salvaged in a GALE-independent manner (49, 50). Our results support some conclusions of prior studies performed in other systems. For example, consistent with the idea that GALE is required to balance NSs, and in agreement with data from galactosemic patients and earlier experimental models (32, 34, 51), we observed a large increase in UDP-Gal in GALE^−/−^ cells in the presence of supplementary Gal (Fig. 1C). On the other hand, our results challenge some conclusions of prior work. For instance, in GALE-deficient human patients and some model systems, Gal is acutely toxic, leading to severe symptoms or impairing cell health and proliferation (34, 52, 53). It has been proposed that the accumulation of intermediate metabolites or toxic byproducts might account for these harmful effects (54). However, we found no evidence of Gal toxicity in our GALE^−/−^ cell systems (Fig. S1). It may be that the accumulation of intermediate metabolites affects only specific cell or tissue types, perhaps due to differences in Gal metabolism. Testing this hypothesis will be an important focus of future studies.

### GALE is required to support glycoconjugate biosynthesis

While aberrant glycosylation has been reported in other subtypes of galactosemia (42, 55– 62), little is known about how loss of GALE affects glycan synthesis in human cells, either in the presence or absence of supplementary Gal. Here, we show that GALE deletion dramatically affects glycoconjugate biosynthesis, with greatly reduced Gal, GalNAc, and sialic acid content in the cell-surface glycans of GALE^−/−^ cells (Fig. 2A), impacting both N- and O-linked glycoproteins (Figs. 2C-F). Moreover, our glycolipid profiling suggests that GALE is essential for the biosynthesis of such species as myelin gangliosides, including lactosylceramide (LacCer) and sulfatides (Fig. 2B). GM3 ganglioside is a regulator of leptin signaling and has been linked to development of insulin resistance (63, 64), sulfatides are major components of myelin and are believed to play crucial roles in neuronal differentiation (65, 66) and deficiencies in LacCer biosynthesis cause locomotor deficits and abnormal brain development in mice (67). Therefore, GALE-dependent glycolipid biosynthesis may be required for normal nutrient responses and neuronal physiology *in vivo*. Our data may also provide a functional explanation for the previously reported postprandial transcriptional upregulation of GALE in multiple tissue types (14, 25), suggesting that GALE could be critical for balancing NS pools and supporting glycoprotein and glycolipid biosynthesis after feeding.

Our results may also have implications for previously unappreciated aspects of human cell metabolism. For example, we observed that Gal supplementation restored the biosynthesis of UDP-GalNAc and GalNAc-bearing glycans in GALE^−/−^ cells (Figs. 1C, 2F). These results were unexpected, because Gal is thought not to enter the well-characterized pathway for UDP-GalNAc biosynthesis (68, 69). We suggest two possible explanations for these observations. First, Gal supplementation may upregulate the activity of salvage enzymes, such as galactokinase (GALK), allowing cells to recycle GalNAc monosaccharides from serum glycoproteins more efficiently. Preliminary experiments suggest that the expression of GALK1 and GALK2 is unaffected by Gal treatment or GALE deletion (Fig. S9), but future studies on GALK enzymatic activity will be needed to test this hypothesis further. Second, very high levels of Gal may result in its non-canonical entry into the hexosamine biosynthetic pathway, which biosynthesizes UDP-GlcNAc from Glc. In this scenario, sufficient UDP-GalNAc may be produced to restore glycoprotein biosynthesis. This hypothesis remains to be tested, but significant substrate promiscuity has been documented previously in mammalian hexosamine metabolism (70–72). Experiments are ongoing to investigate these two, mutually compatible possibilities.

### GALE loss triggers receptor hypoglycosylation and dysfunction

Our results demonstrate that GALE is required to support cell surface receptor signaling even in nutrient-replete human cells, with significant effects on death receptor function (Figs. 3 and S8). Aberrant glycosylation has long been known to affect apoptotic receptor signaling. For example, oncogene activation upregulates sialyltransferase expression and glycoprotein sialylation in colon adenocarcinomas, promoting cell migration and resistance to galectin-mediated apoptosis (73–75). Similarly, hypersialylation of Fas by ST6Gal-I protects tumor cells from Fas-mediated apoptosis (48). Given the dramatic loss of global sialylation in GALE^−/−^ cells (Figs. 2A-E), we propose that Fas hyposialylation accounts for their hypersensitivity to FasL-induced apoptosis (Figs. 3C and S8). Consistent with this notion, we observed GALE-dependent differences in FasL-mediated caspase-3 activation (Fig. 3D). Importantly, however, these results are not attributable to a general predisposition to apoptosis, because no such hypersensitivity was observed in response to staurosporine (Fig. S7). Beyond Fas, we also observed aberrant glycosylation of a subset of integrins in GALE^−/−^ cells (Figs. 3B and S5). Glycosylation is well known to impact the function of several integrins, particularly β_1_ (74, 76–83), and a very recent study suggested that GALE, in particular, may be necessary for integrin β_1_ function in platelet homeostasis (29). Based on these studies and our own results, we speculate that GALE activity may be required in some contexts for normal integrin-mediated attachment to extracellular matrix, cell adhesion and migration, and tissue homeostasis. Experiments are underway to test this hypothesis.

Finally, our results may have implications for understanding global protein N-glycosylation in both model systems and galactosemic patients. Past studies of mammalian GALE have largely focused on its role in O-glycan biosynthesis (35, 84–86). Therefore, it is especially noteworthy that we discovered alterations in the N-glycans of several cell surface receptors (Figs. 3 and S5). Altered N-glycans have been implicated in other subtypes of galactosemia, based on observations in both human patients (42) and animal models (55). However, the role for mammalian GALE in supporting N-glycan biosynthesis has received little attention in any clinical or experimental context. We propose that altered N-glycans may contribute to the pathology of GALE-deficient galactosemia, and GALE may play a critical role in healthy tissue under normal conditions that trigger glycosylation changes, such as feeding or stress (14, 15).

### Conclusion

NS metabolic enzymes have been extensively characterized at the biochemical level. However, the mechanisms and functions of dynamic changes in NS pools in cells and organisms remain poorly understood. We have deleted GALE, a key regulator of NS levels, in human cell systems as a model for NS dysregulation. Our data demonstrate that human GALE is required for normal glycoconjugate biosynthesis and receptor signaling, even in nutrient-replete cells. Moreover, we anticipate that the dramatic absence of cell surface Gal, GalNAc and sialic acid in GALE^−/−^ cells, combined with the ability to restore normal glycosylation with simple Gal supplementation, will make GALE^−/−^ cells an attractive model system for studying glycan function in normal cell physiology, metabolic syndrome, thrombocytopenia and galactosemia.

## Experimental procedures

### Cell culture

293T and HeLa cells were cultured in Dulbecco’s Modified Eagle’s Medium (DMEM) containing 10% fetal bovine serum (FBS), 100 g/ml streptomycin, and 100 units/ml penicillin in 5% CO_2_ at 37 °C. For monosaccharide supplementation experiments, cells were preconditioned with 250 µM mannitol or Gal (Sigma-Aldrich) for 72 hours, with replenishment of monosaccharide every 24 hours.

### Generation of GALE^−/−^ cell lines

GALE^−/−^ HeLa and 293T cell lines were constructed essentially as described (87). Briefly, cells at ~50% confluency were stably transduced with LentiCas9 virus obtained from the Duke Functional Genomics Facility in the presence of 4 µg/ml Polybrene. After overnight incubation, medium was replaced and cells were allowed to recover for 48 hours before selection with 3 (HeLa) or 5 (293T) µg/ml blasticidin. Cells were passaged until an uninfected control plate had no live cells remaining. Following selection, cells were infected with lentiviruses bearing one of three single-guide RNA (sgRNA) sequences targeting the GALE coding sequence or a “safe harbor” AAVS1-targeting control sgRNA (88). After sgRNA infection, cells were selected for stable sgRNA expression with 1.5 µg/ml (HeLa) or 0.5 µg/ml (293T) puromycin, with continued presence of blasticidin. Following drug selection, clonal lines were generated via limiting dilution and assayed for GALE deletion via Western blot.

The GALE sgRNA sequences used were: GALE-1: GAGAAGGTGCTGGTAACAGG GALE-2: GGAGGCTGGCTACTTGCCTG GALE-3: GCCAGGTGCCATGGCAGAGA

### Western blotting

Protein samples were quantified by bicinchoninic acid (BCA) assay according to the manufacturer’s protocol (ThermoFisher). Samples with equal protein amounts were separated by sodium dodecyl sulfate polyacrylamide gel electrophoresis (SDS-PAGE) using standard methods (89) and electroblotted onto polyvinylidene fluoride (PVDF) membrane (ThermoFisher, 88518). After blocking in Tris-buffered saline with 0.1% Tween (TBST) and 2.5% dry milk, blots were incubated overnight at 4 °C in primary antibody with 5% bovine serum albumin (BSA) in TBST. The following day, blots were washed three times in TBST, incubated with an appropriate horseradish peroxidase-conjugated secondary antibody (Southern Biotech) for 1 hour at room temperature, washed three times in TBST, and developed via enhanced chemiluminescence (ECL) according to the manufacturer’s instructions (WesternBright ECL, Advansta). The following primary antibodies were used: GALE (Abcam ab 118033), tubulin (Sigma-Aldrich T6074), Fas (Cell Signaling C18C12), caspase-3 (Cell Signaling 8G10), FADD (Abclonal A5819), integrin α_5_ (Cell Signaling 4705), integrin β_1_ (Santa Cruz sc-13590), integrin β_5_ (Cell Signaling D24A5) and GAPDH (Cell Signaling 14C10).

### Protein N-glycosidase-F digestion

Cells were harvested in 135 mM KCl/15 mM sodium citrate with 10 minutes of gentle rocking, washed in phosphate-buffered saline (PBS) and resuspended in lysis buffer (20 mM Tris-HCl, 150 mM NaCl, 1% Triton-X 100, pH 7.5). N-glycans were digested using PNGase-F (New England Biolabs, P0704S) essentially according to manufacturer’s protocol but with a three-hour incubation at 37 °C.

### Flow cytometry

For both lectin- and antibody-labeled flow cytometry, cells were harvested in 135 mM KCl/15 mM sodium citrate with 10 minutes of gentle rocking and resuspended in PBS with 2% BSA at 1 million cells/ml. For lectin labeling, cells were rotated for 30 minutes at 4 °C with fluorescent lectin in PBS/BSA, washed three times with PBS/BSA, and fixed with 2% paraformaldehyde for 20 minutes before analysis on a FACSCanto II cytometer (BD biosciences). The lectins used were fluorescein-SNA (vector labs FL-1301-2), fluorescein-WGA (vector labs FL-1201) and fluorescein-jacalin (vector labs FL-1121).

For antibody labeling, aliquots of cells were incubated with primary antibody for 30 minutes at 4 °C and then washed three times with PBS/BSA. Then, cells were washed twice with 2% BSA/PBS and incubated with secondary antibody for 30 minutes. After the final incubation, cells were fixed with 2% paraformaldehyde for 20 minutes before analysis on a FACSCanto II cytometer (BD biosciences). The antibodies used were Fas (Biolegend 305602) and Goat anti-mouse Alexa Fluor 488 (Thermo A11001).

### Glycolipid profiling

Lipid extraction was performed using a modified Bligh-Dyer method (90). Briefly, cell pellets were resuspended in 1.6 ml PBS and transferred into a 17 ml glass tube with a Teflon-lined cap. Them, 4 ml methanol and 2 ml chloroform were added to create a single-phase Bligh-Dyer solution (chloroform:methanol:PBS 1:2:0.8). The mixture was vigorously vortexed for 2 minutes and then sonicated in a water bath at room temperature for 20 minutes. After centrifugation at 3,000 g for 10 minutes, supernatants were transferred to fresh glass tubes and acidified by adding 100 μl 37% HCl. After mixing, acidified solutions were converted into two-phase Bligh-Dyer systems by adding 2 ml PBS and 2 ml chloroform. After centrifugation at 3,000 g for 10 minutes, the lower phase was collected and dried under nitrogen. Lipid extracts were stored at −20 °C until LC/MS analysis. For lipidomic analysis, samples were dissolved in a 200 μl solution of chloroform:methanol (2:1) and 20 μl were injected for normal-phase LC/MS analysis.

Normal-phase LC was performed on an Agilent 1200 Quaternary LC system equipped with an Ascentis silica high performance liquid chromatography (HPLC) column (5 μm, 25 cm x 2.1 mm, Sigma-Aldrich). Mobile phase A was chloroform:methanol:aqueous NH_4_OH (800:195:5, by volume); mobile phase B was chloroform:methanol:water:aqueous NH_4_OH (600:340:50:5, by volume); mobile phase C was chloroform:methanol:water:aqueous NH_4_OH (450:450:95:5, by volume). The elution was performed as follows: 100% mobile phase A was held isocratically for 2 minutes, then linearly increased to 100% mobile phase B over 14 minutes and held at 100% B for 11 minutes. The gradient was then changed to 100% mobile phase C over 3 minutes and held at 100% C for 3 minutes, and finally returned to 100% A over 0.5 minutes and held at 100% A for 5 minutes. The LC eluent (total flow rate of 300 μl/minute) was introduced into the electrospray ionization (ESI) source of a high resolution TripleTOF 5600 mass spectrometer (Sciex). Instrument settings for negative ion ESI and MS/MS analysis of lipid species were as follows: IS = −4500 V; CUR = 20 psi; GSI = 20 psi; DP = −55 V; and FP = −150 V. MS/MS analysis used nitrogen as the collision gas. Data analysis was performed using Analyst TF1.5 software (Sciex).

### Glycan profiling

Frozen cell pellets were resuspended in methanol:water (4:1.5) and sheared using a 20-gauge needle-equipped syringe. Delipidation was achieved by adding chloroform to a final ratio of 2:4:1.5 chloroform:methanol:water. Samples were incubated for 2 hours, centrifuged, decanted and incubated in the same chloroform/methanol/water mixture overnight. The next day, samples were centrifuged, resuspended in 4:1 acetone:water and incubated on ice for 15 minutes. Samples were centrifuged and supernatant was decanted and incubated with acetone:water once more. De-lipidated protein was dried under nitrogen and stored at −20 °C until analysis.

For N- and O-glycan analysis, 2-4 mg of dried protein powder was used per sample. N- and O-glycans were released enzymatically or chemically, respectively, and isolated as previously described (91). Following clean-up, glycans were permethylated as previously described (92), dried under nitrogen and stored at −20 °C until analysis.

Permethylated glycans were resuspended and mixed with an internal standard before analysis via reversed-phase LC-MS/MS on a Thermo Scientific Velos Pro mass spectrometer as described (91). Glycan structures were manually interpreted based on in-house fragmentation rules, augmented by GlycoWorkbench and GRITS Toolbox semi-automated software solutions (93, 94). All glycan structures are depicted using standard symbol nomenclature for glycobiology (95).

### Glycan monosaccharide analysis

Complete hydrolysis of isolated N- and O-glycans to monosaccharides was performed using 2 M trifluoroacetic acid (TFA) prior to analysis by HPAEC with pulsed amperometric detection (PAD). 100 µg of dry glycan sample was dissolved in 50 µl of ultrapure water and 50 μl 4 M TFA and mixed thoroughly. Samples were hydrolyzed at 100 °C for 4 hours. Hydrolyzed samples were cooled to room temperature and then centrifuged at 2,000 rpm for 2 minutes. Acid was removed by complete evaporation under dry nitrogen flush, followed by two rounds of co-evaporation with 100 µl 50% isopropyl alcohol each time. Dry samples were resuspended in 142 µl ultrapure water and 70 µg sample was injected for HPAEC-PAD analysis.

For neutral and amino sugar analysis, HPAEC-PAD profiling was performed on a Dionex ICS-3000 system equipped with a CarboPac PA1 column (4 × 250 mm), guard column (4 × 50 mm) and PAD using standard Quad waveform supplied by the manufacturer. An isocratic mixture of 19 mM NaOH with 0.95 mM NaOAc was used at a flow rate of 1 ml/minute for 20 minutes. Neutral and amino sugars were identified and quantified by comparison with an authentic standard mixture of L-fucose, D-galactosamine, D-glucosamine, D-galactose, D-glucose and D-mannose using Thermo Scientific Chromeleon software (version 6.8).

For sialic acid analysis, 50 μg of samples were hydrolyzed at 80 °C using 2 M acetic acid for 3 hours. Released sialic acids were isolated by spin-filtration using a 3,000 kDa molecular weight cutoff centrifugal device (Nanosep 3K Omega, PALL Life Sciences, OD003C34). The flow-through containing released sialic acid was dried and derivatized using 1,2-diamino-4,5-methylene-dioxybenzene (DMB) (Sigma-Aldrich, D4784-10MG), as previously described (96). Fluorescent, DMB-derivatized sialic acids were analyzed by reversed-phase HPLC using a Thermo-Dionex UltiMate 3000 system equipped with a fluorescence detector. Samples were isocratically eluted with 9% acetonitrile (9%) and 7% methanol in ultrapure water over 30 minutes using Acclaim 120-C18 column (4.6 × 250 mm, Dionex) at a flow rate of 0.9 ml/minute. The excitation and emission wavelengths were set at 373 nm and 448 nm, respectively. DMB-derivatized sialic acids were identified and quantified by comparing the elution times and peak areas to known authentic standards of *N*-acetylneuraminic acid and *N*-glycolylneuraminic acid using Chromeleon software (version 6.8).

### Cell viability assays

Cells were plated into 96-well plates at 5,000 cells/well in phenol red-free DMEM and allowed to recover for 24 hours. FasL (Adipogen AG-40B-0130-C010) or staurosporine (LC labs S-9300) was added and incubated for the indicated times. Then, cell viability was assessed by adding soluble formazan (MTS assay, Promega) and incubating for 1 hour at 37 °C, 5% CO_2_, before measuring absorbance at 490 nm.

### NS analysis

Cells were cultured to confluency in 10 cm culture dishes, detached using 0.25% trypsin at 37 °C and washed three times with PBS at 4 °C. Cells were lysed in methanol on dry ice, vigorously vortexed and centrifuged to pellet proteinaceous solids. Pellets were resuspended in 8 M urea and total protein was quantified by BCA assay. Methanol supernatants (containing NSs) were SpeedVac-dried and resuspended in 80 mM Tris-HCl pH 7.4. Samples were separated on a Dionex CarboPac PA1 column (Thermo) using a Dionex ICS-5000+ SP HPAEC instrument and photodiode array. 10 µl of each sample was injected onto the column after pre-equilibration with 1 mM NaOH (solvent A). Samples were eluted at a flow rate of 1 ml/min using solvent A and solvent B (1 mM NaOH, 1M NaOAc) as follows: 0-100% B for 0-28 minutes, 100% B for 2 minutes. Samples were quantified using absorbance at 260 nm and compared to a series of authentic UDP-Gal or UDP-GalNAc standards at known concentrations (Sigma-Aldrich). Peak areas were integrated using Chromeleon software and cellular NS concentrations were calculated using standard curves and total protein amounts.

### Quantification and Statistical Analysis

No randomization was performed for these studies. Quantitative experiments (e.g., HPAEC analysis, MTS viability assays) were performed with a minimum of three biological replicates originating from independent cultures. The number of biological replicates and statistical tests used are given in the figure legends. All Western blots are representative of at least two experiments from biologically independent samples.

## Acknowledgements

We thank So Young Kim and the Duke Functional Genomics Facility staff for designing and testing sgRNAs, Abhi Chhetri and Kenichi Yokoyama for training and assistance with HPAEC NS quantification, Biswa Choudhury and the UCSD Glycotechnology core facility for monosaccharide composition analysis and Boyce Lab members for helpful discussion. This work was supported by a Scholar Award from the Rita Allen Foundation (to M.B.), NIH grants GM069338 and EY023666 (to Z.G.), Welch Foundation grant I-1686 (to J.J.K.) and a postdoctoral fellowship from the German Academic Exchange Service (to N.N.). L.W. is co-director of the Thermo fisher appointed Center of Excellence in Glycoproteinics at the CCRC.

---

**Figure S1.**
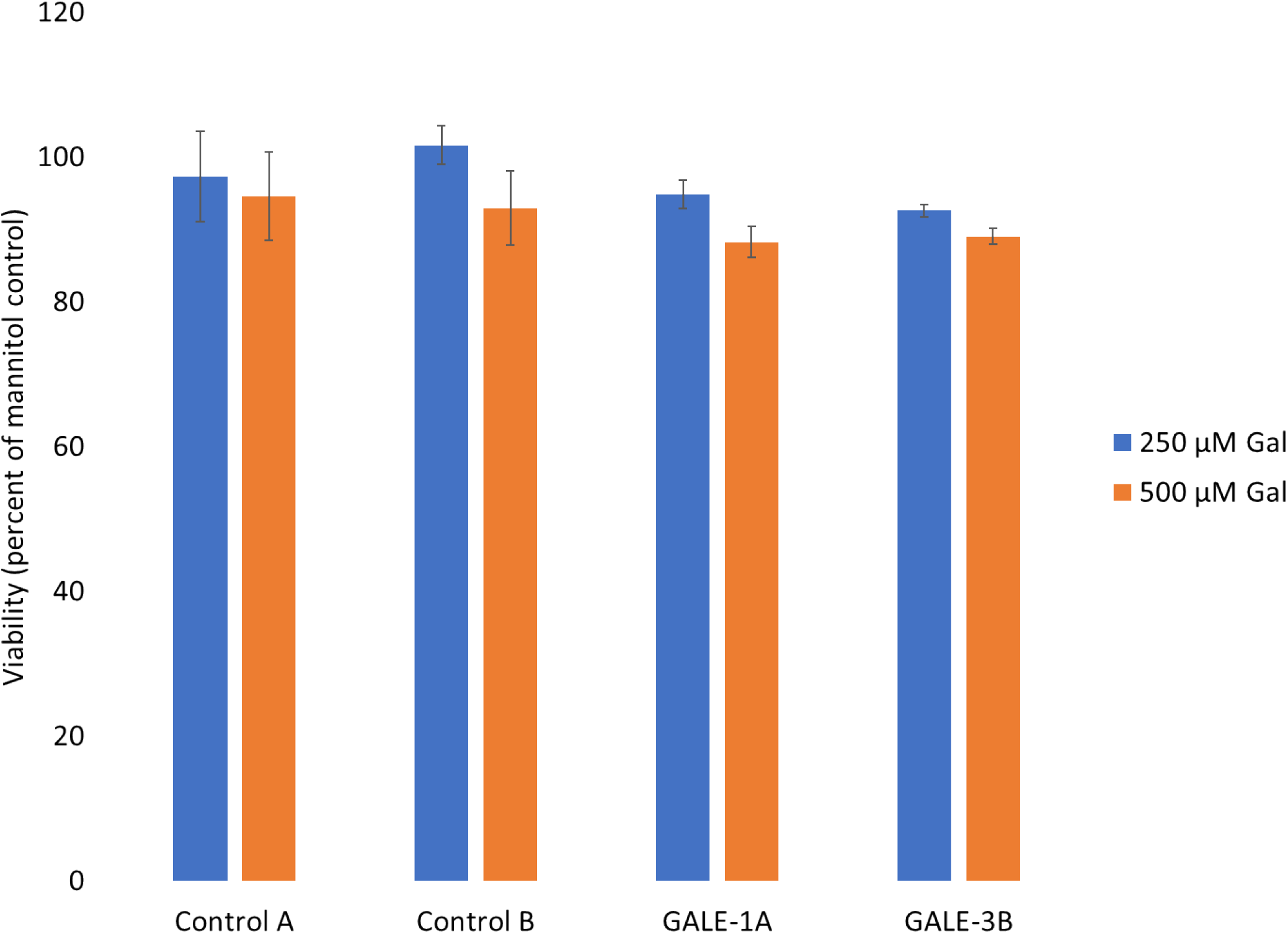
GALE deletion does not affect cell viability. Related to Figure 1. Control and GALE^−/−^ HeLa cell clones were treated with 250 or 500 μM Gal or mannitol for 72 hours. Cell viability was measured by MTS assay and normalized to mannitol controls. N=3 biological replicates. Error bars represent SEM. No statistically significant difference exists between control and GALE^−/−^ cells (two-way ANOVA).

**Figure S2.**
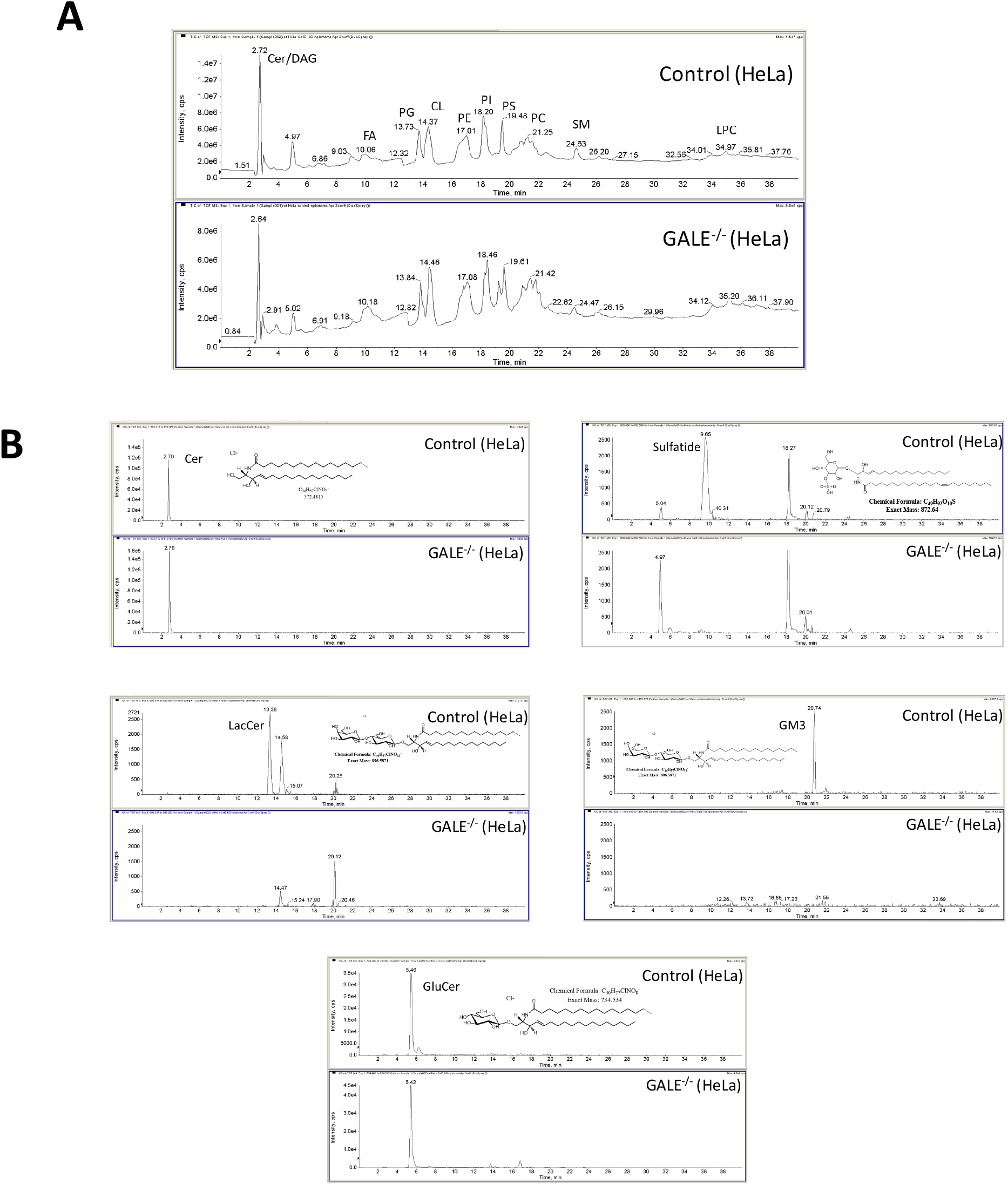
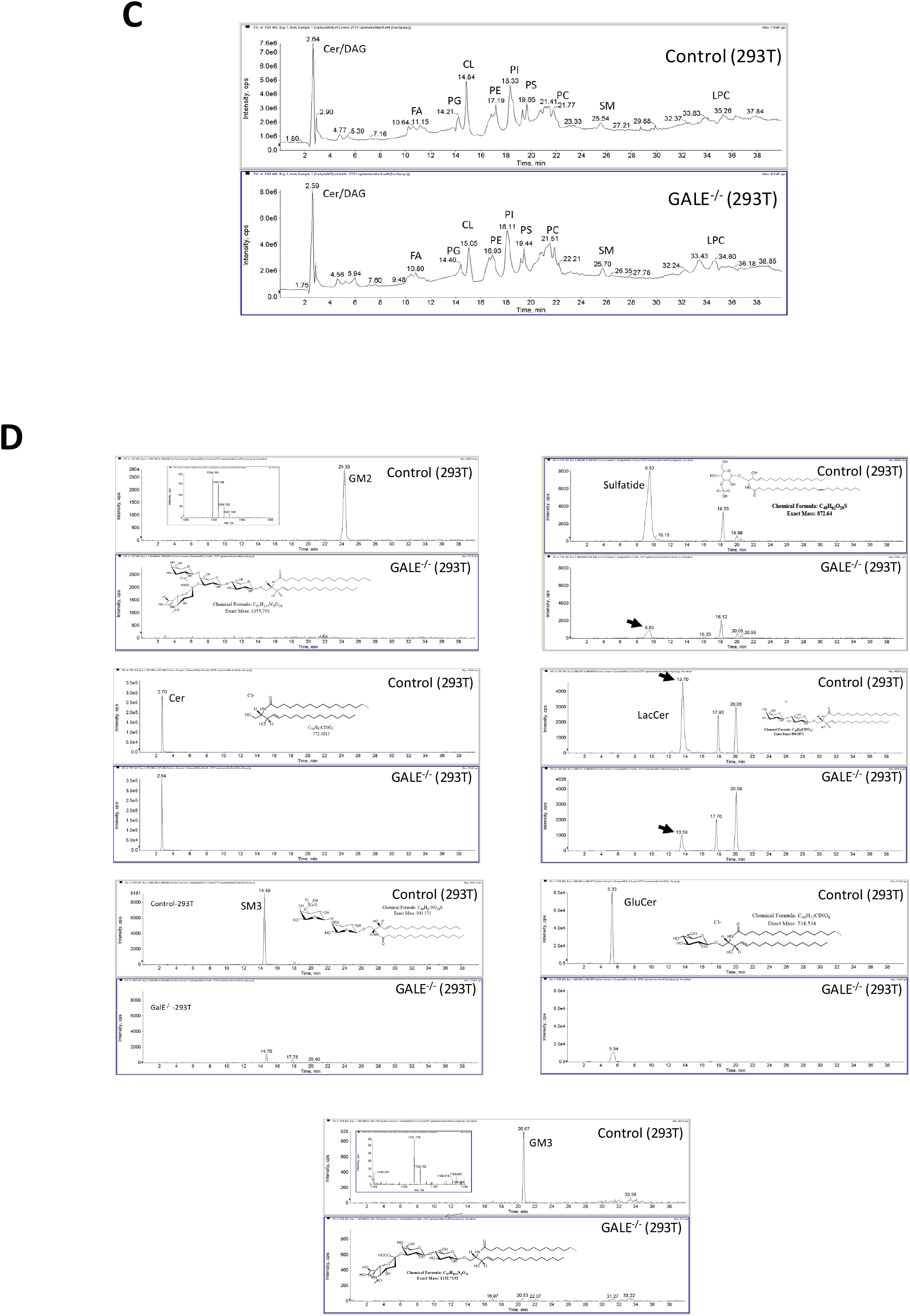
Normal phase LC-MS analysis of total lipid extracts from control and GALE^−/−^ cells. Related to Figure 2. Lipids were isolated from control and GALE^−/−^ HeLa and 293T cells and analyzed by LC-MS. Total (A) and extracted (B) ion chromatograms from control and GALE^−/−^ HeLa cells. Total (C) and extracted (D) ion chromatograms from control and GALE^−/−^ 293T cells.

**Figure S3.**
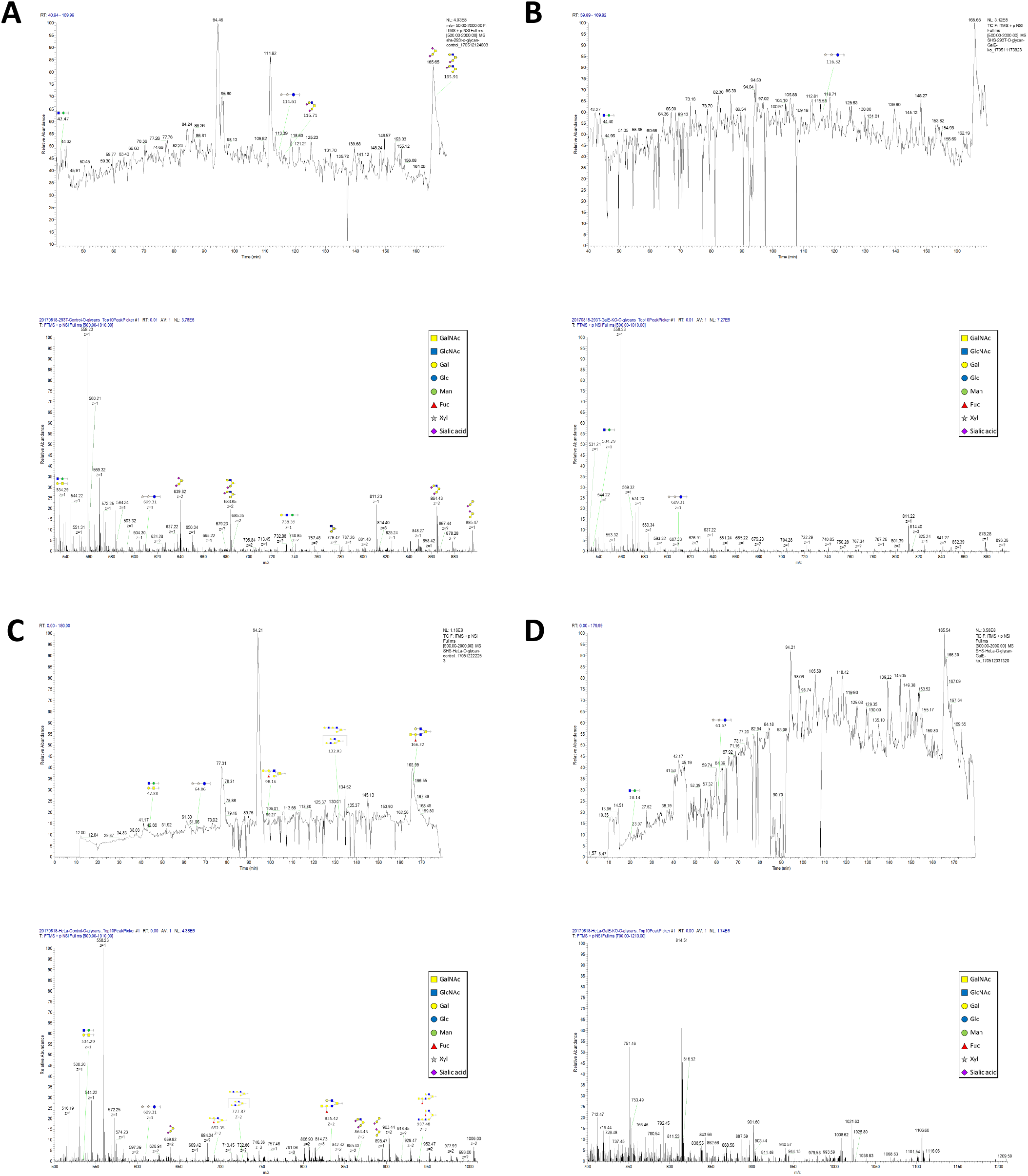
Captured ion chromatograms from O-linked glycan analysis of control and GALE^−/−^ cells. Related to Figure 2. O-glycans from control 293T (A), GALE^−/−^ 293T (B), control HeLa (C) and GALE^−/−^ HeLa (D) cells were analyzed by LC-MS.

**Figure S4.**
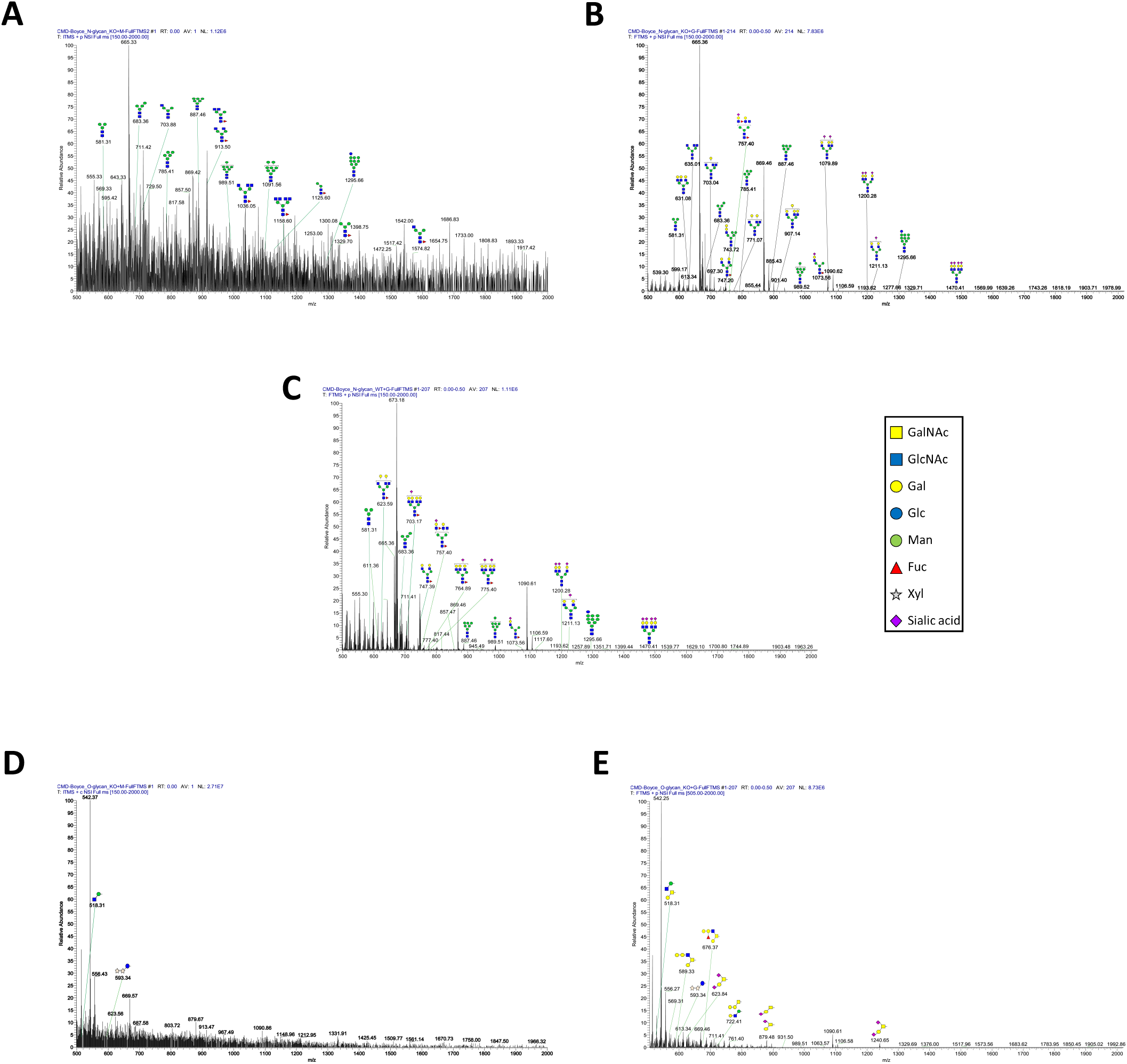
Captured ion chromatograms from glycan analysis of GALE^−/−^ HeLa cells, treated with or without Gal. Related to Figure 2. Control or GALE^−/−^ HeLa cells were treated with 250 µM Gal or mannitol for 72 hours and glycans were analyzed by LC-MS. (A) N-glycans from mannitol-treated GALE^−/−^ cells. (B) N-glycans from Gal-treated GALE^−/−^ cells. (C) N-glycans from Gal-treated control cells. (D) O-glycans from mannitol-treated GALE^−/−^ cells. (E) O-glycans from Gal-treated GALE^−/−^ cells.

**Figure S5.**
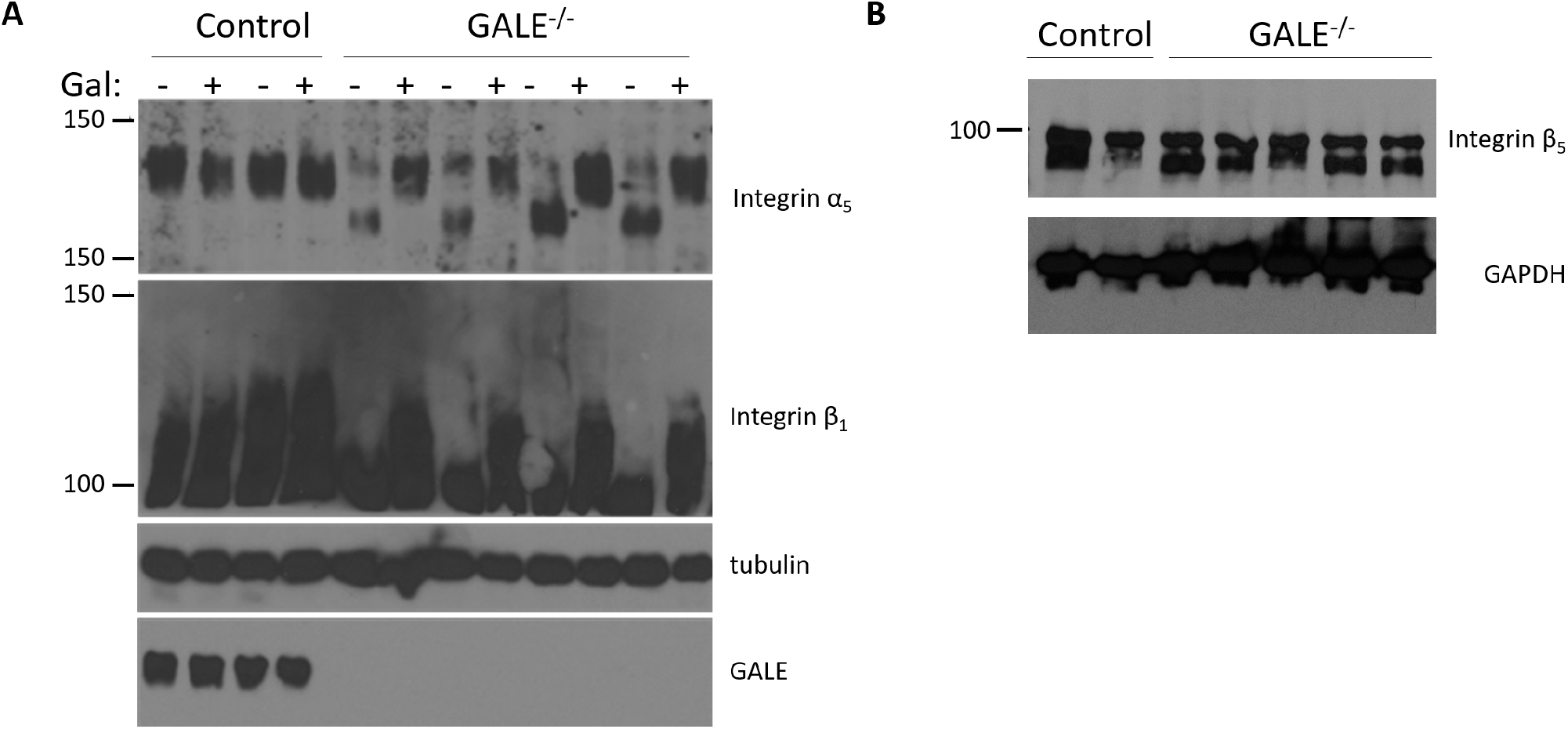
GALE is required for the glycosylation of a subset of integrins. Related to Figure 3. Control or GALE^−/−^ HeLa clones were treated with 250 µM Gal or mannitol for 72 hours and lysates were analyzed by Western blot. Integrins α_5_ and β_1_ (A), but not β_5_ (B), are hypoglycosylated in GALE-/-cells.

**Figure S6.**
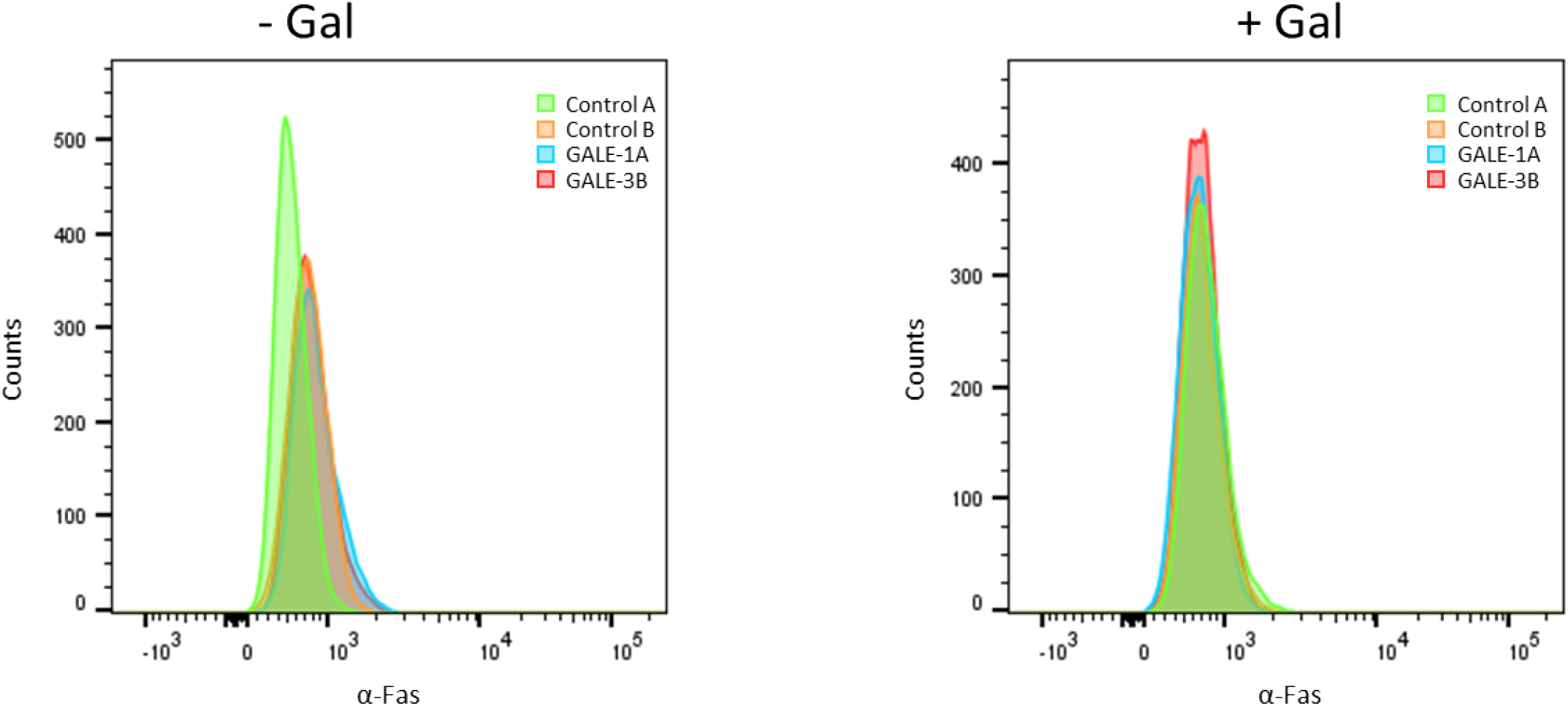
Fas cell surface expression is not affected by GALE deletion or Gal supplementation. Related to Figure 3. Control or GALE^−/−^ HeLa clones were treated with 250 µM Gal or mannitol for 72 hours, stained with an anti-Fas antibody and analyzed by flow cytometry.

**Figure S7.**
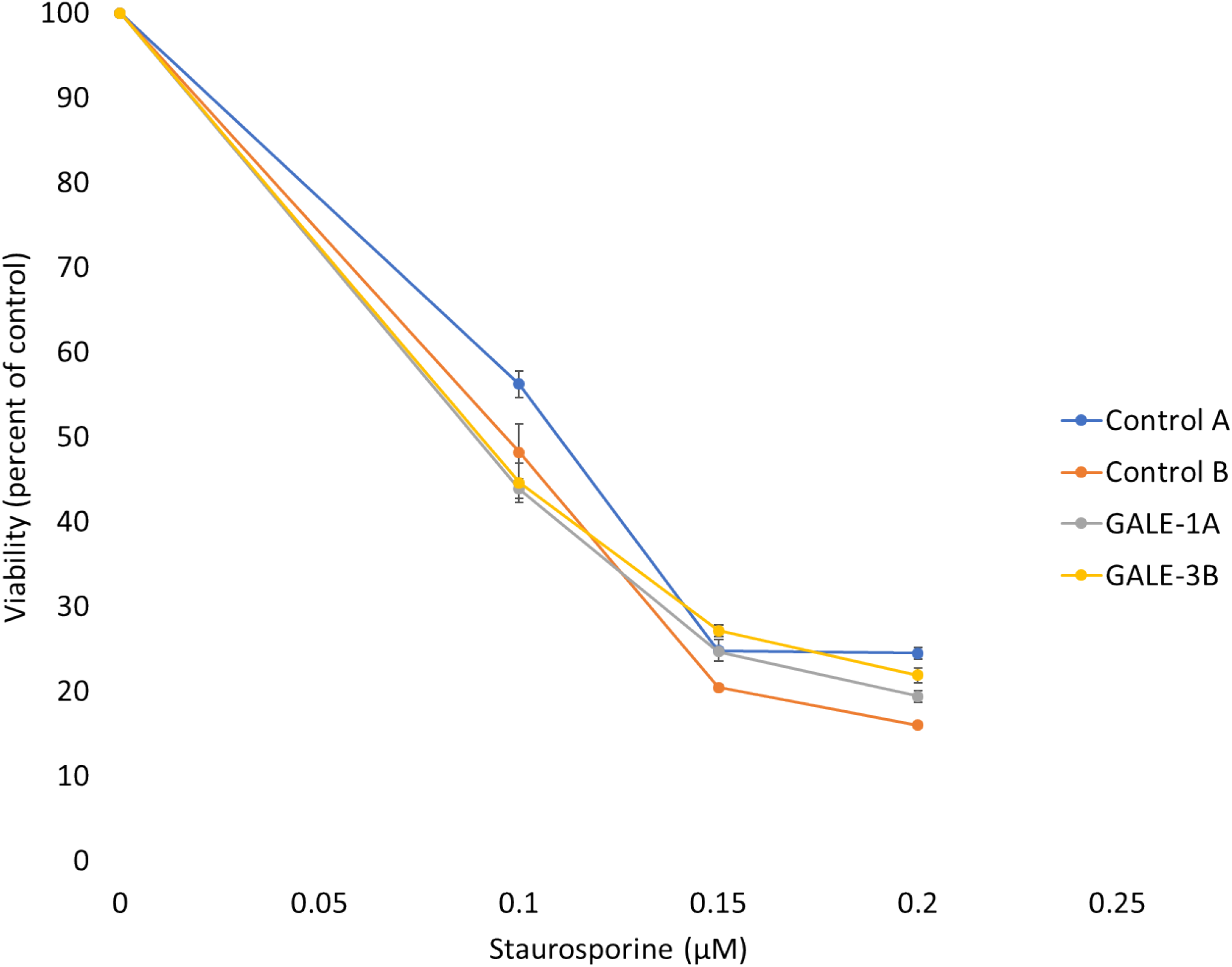
GALE deletion does not generally sensitize cells to apoptosis. Related to Figure 3. Control or GALE^−/−^ HeLa clones were treated with the indicated concentrations of staurosporine for 24 hours and cell viability was measured by MTS assay. N=3 biological replicates. Error bars represent SEM.

**Figure S8:**
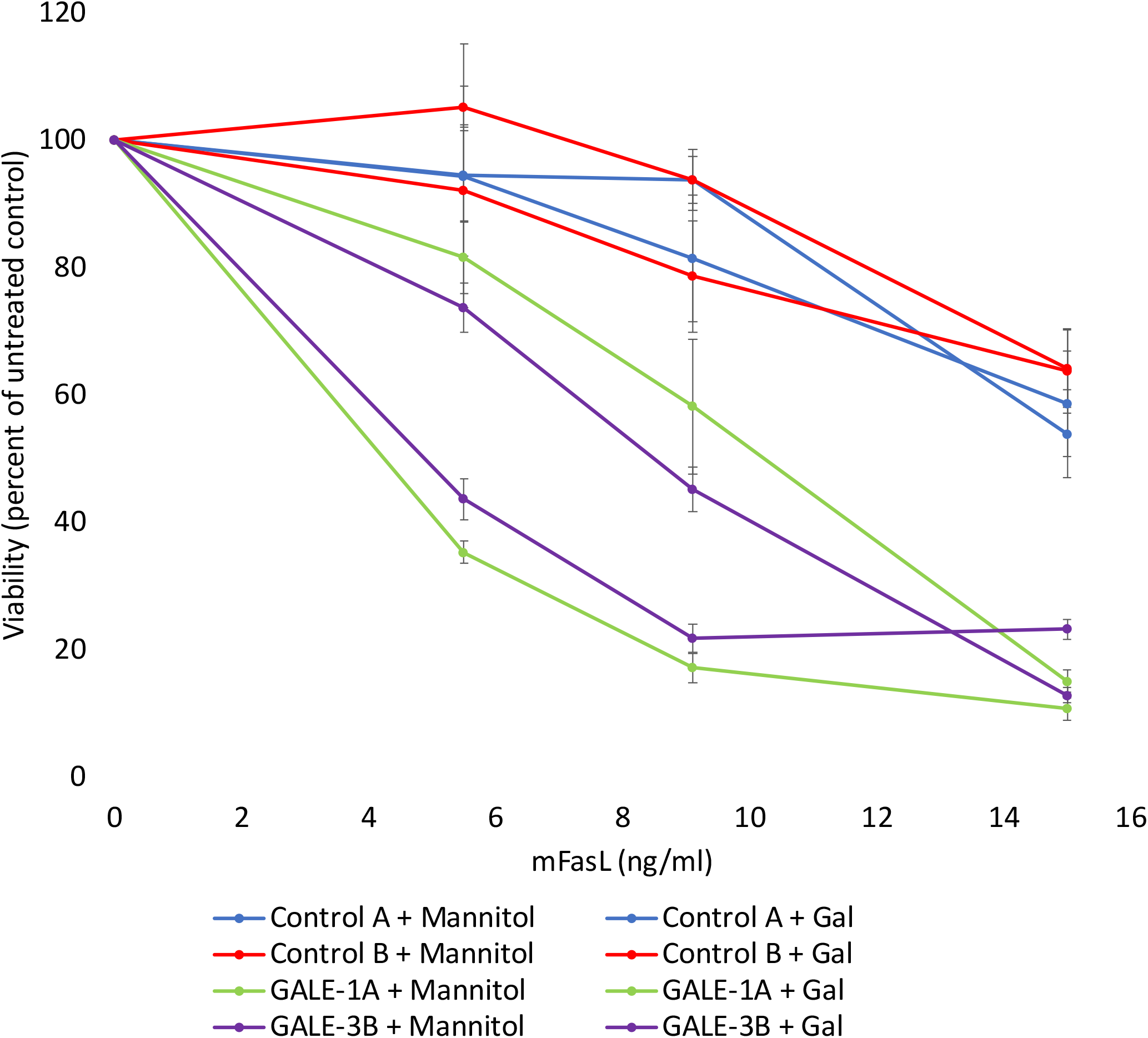
Gal supplementation reverts the hypersensitivity of GALE^−/−^ cells to FasL. Related to Figure 3. Control or GALE^−/−^ HeLa clones were treated with 250 µM mannitol or Gal for 72 hours and then with the indicated concentrations of FasL for an additional 24 hours. Cell viability was measured by MTS assay. N=3 biological replicates. Error bars represent SEM. *p* < 0.05 comparing mannitol-versus Gal-treated GALE^−/−^ cells at 5.5 ng/ml and 9.1 ng/ml FasL (three-way ANOVA, post hoc Tukey’s HSD test).

**Figure S9:**
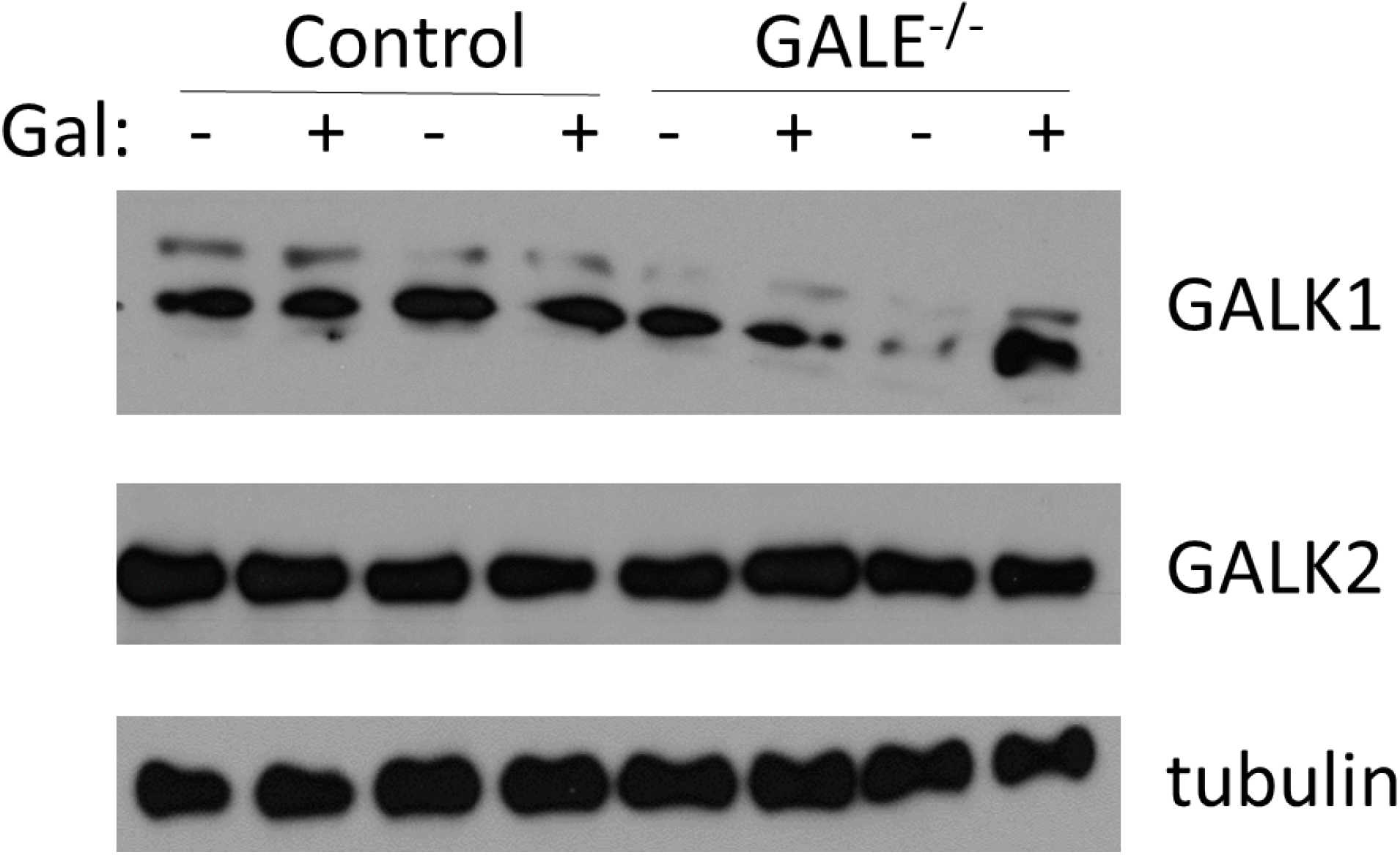
Expression of GALK1 and GALK2 is not affected by GALE deletion or Gal supplementation. Related to Figures 1 and 2. Control or GALE^−/−^ HeLa clones were treated with 250 µM Gal or mannitol for 72 hours and lysates were analyzed by Western blot.

